# Population genomics of hypervirulent *Klebsiella pneumoniae* clonal group 23 reveals early emergence and rapid global dissemination

**DOI:** 10.1101/225359

**Authors:** Margaret MC Lam, Kelly L Wyres, Sebastian Duchêne, Ryan R Wick, Louise M Judd, Yunn-Hwen Gan, Chu-Han Hoh, Sophia Achuleta, James S Molton, Shirin Kalimuddin, Tse Hsien Koh, Virginie Passet, Sylvain Brisse, Kathryn E Holt

## Abstract

Since the mid-1980s there have been increasing reports of severe community-acquired pyogenic liver abscess, meningitis and bloodstream infections caused by hypervirulent *Klebsiella pneumoniae*, predominantly encompassing clonal group (CG) 23 serotype K1 strains. Common features of CG23 include a virulence plasmid associated with iron scavenging and hypermucoidy, and a chromosomal integrative and conjugative element (ICE) encoding the siderophore yersiniabactin and the genotoxin colibactin. Here we investigate the evolutionary history and genomic diversity of CG23 based on comparative analysis of 98 genomes. Contrary to previous reports with more limited samples, we show that CG23 comprises several deep branching sublineages dating back to the 1870s, many of which are associated with distinct chromosomal insertions of ICEs encoding yersiniabactin. We find that most liver abscess isolates (>80%) belong to a dominant sublineage, CG23-I, which emerged in the 1920s following acquisition of ICE*Kp10* (encoding colibactin in addition to yersiniabactin) and has undergone clonal expansion and global dissemination within the human population. The unique genomic feature of CG23-I is the production of colibactin, which has been reported previously as a promoter of gut colonisation and dissemination to the liver and brain in a mouse model of CG23 *K. pneumoniae* infection, and has been linked to colorectal cancer. We also identify an antibiotic-resistant subclade of CG23-I associated with sexually-transmitted infections in horses dating back to the 1980s. These data show that hypervirulent CG23 *K. pneumoniae* was circulating in humans for decades before the liver abscess epidemic was first recognised, and has the capacity to acquire and maintain AMR plasmids. These data provide a framework for future epidemiological and experimental studies of hypervirulent *K. pneumoniae*. To further support such studies we present an open access and completely sequenced human liver abscess isolate, SGH10, which is typical of the globally disseminated CG23-I sublineage.

## Introduction

*Klebsiella pneumoniae* (*Kp*) is a ubiquitous bacterium and important cause of multidrug-resistant healthcare-associated infections. The past three decades have also seen the emergence of severe community-acquired hypervirulent *Kp* disease, usually manifesting as pyogenic liver abscess with accompanying bacteraemia, but also meningitis, brain abscess or opthalmitis ^1,2^. Earliest reports of this syndrome emerged across parts of Asia including Taiwan, China, Hong-Kong, Singapore and South Korea ^3-6^. More recently, reports have emerged from Europe, the United States, South America, the Middle East and Australia ^7-13^.

The majority of *Kp* liver abscess isolates belong to a small number of clonal groups (CGs) ^14-17^ defined by multi-locus sequence typing (MLST) ^18^. In particular, CG23 was shown to account for 37-64% isolates in Taiwan ^16^, Singapore ^19^ and mainland China ^15^,^20^. CG23 *Kp* are associated with the highly serum-resistant K1 capsule and a number of virulence factors; the genotoxin colibactin, microcin E492, and the iron-scavenging siderophores aerobactin, yersiniabactin and salmochelin ^1,21,22^. The siderophores are associated with the ability to cause disseminated infection in mouse models ^22-25^ and with invasive infections in humans ^21,25,26^.

Two CG23 comparative genomic analyses have been published, incorporating up to 27 isolates ^27,28^. Both studies reported limited nucleotide variation in core chromosomal genes and high conservation of the acquired virulence genes. These include the K1 capsule synthesis locus KL1, the chromosomally-encoded yersiniabactin locus (*ybt*), plus the *iro* (salmochelin), *iuc* (aerobactin) and *rmpA/rmpA2* genes (upregulators of capsule expression) located on the pK2044 virulence plasmid ^29^. The *ybt* locus is mobilised by the ICE*Kp* integrative conjugative element ^30^, for which we have recently described over a dozen variants in the wider *Kp* population ^26^. Variants harbouring *ybt,* plus *iro* (ICE*Kpl*) or the colibactin locus (*clb*, ICE*Kp10*) have been reported in CG23 ^26,30,31^. ICE*Kpl* was the first ICE to be described in *Kp* ^30^, originating from the sequence type (ST) 23 liver abscess strain NTUH-K2044 which was also the first CG23 strain to be completely sequenced ^29^. However ICE*Kp10* is more common and was observed in all other CG23 genomes investigated in Struve *et al* ^28^.

CG23 has not generally been associated with acquired antimicrobial resistance (AMR), but the last few years have seen increasing reports of resistant strains, including those resistant to third generation cephalosporins and carbapenems^27,32-34^. The potent combination of virulence and AMR determinants could make these strains a substantial public health threat, but it is not yet clear how often they emerge or whether they can disseminate. To fully evaluate the threat we require a clear understanding of the history of this clone.

Here we describe an updated evolutionary history for CG23 based on genomic analysis of 98 human and equine associated isolates, representing the largest and most geographically diverse collection to date. Our data reveal previously undetected population structure, provide sufficient temporal signal to estimate the date of emergence of CG23, and indicate that the commonly used NTUH-K2044 reference strain is not representative of a ‘typical’ CG23.

## Results

### Phylogenomics and evolutionary history of CG23

Strain CAS686 was previously reported to be a recombinant hybrid strain of ST23 ^28^. Here we identified the likely hybridisation partner to be a ST281-related *Kp* (see **Supplementary Results** and **Fig. S1**). As approximately half of the CAS686 genome comprises non-CG23 DNA, we excluded it from the main CG23 analysis. Phylogenomic analysis of the remaining 97 CG23 genomes (**Table S1**) identified no further recombination events, and showed they were highly conserved across core chromosomal genes, with a median pairwise distance of 233 SNPs (range 1-444 SNPs), and 0.0045% nucleotide divergence (range 0.000019% -0.0086%).

Bayesian and maximum likelihood (ML) phylogenetic analyses indicated that CG23 has a number of deep-branching sublineages, one of which (labelled CG23 sublineage I, CG23-I) has expanded and become globally distributed (**Fig. 1, Fig. S2**). The CG23-I clade was strongly supported by both analysis methods (>99% posterior support in Bayesian tree, 100% bootstrap support in ML tree) and was separated from the rest of CG23 by 49 SNPs, including one nonsense mutation (**Table S2** and **Supplementary Results**). The nonsense mutation is predicted to truncate the outer membrane usher domain of KpcC, likely preventing expression of the Kpc fimbriae, which were discovered in the NTUH-K2044 genome and have been proposed to be characteristic of K1 liver abscess strains ^28,35^. CG23-I comprised 81 isolates (83.5% of all CG23) collected from Asia, Australia, North America, Europe and Africa (**Fig. 1**), including 82% of all liver abscess strains (**Table S1**). *Kp* is considered an important cause of sexually transmitted disease in horses ^36^ but little is known about the molecular epidemiology of this group. The horse isolates included in this study (including genital tract, sperm, foetus and metritis specimens isolated between 1980 and 2004) were selected only on the basis of K1 serotype, but formed a single subclade within CG23-I (**Fig. 1**), separated from the rest of CG23-I by 83 SNPs (**Table S3** and **Supplementary Results**). The oldest CG23 strains M109 (1932, Murray Collection ^37^) and NCTC9494 (1954, human sputum) were both deep branching in the CG23 tree and were not part of CG23-I (**Fig. 1**). Neither was NTUH-K2044, the first sequenced ST23 strain that has served as a reference for much of the reported experimental and genomic work on ST23 ^28-30^.

**Figure 1:**
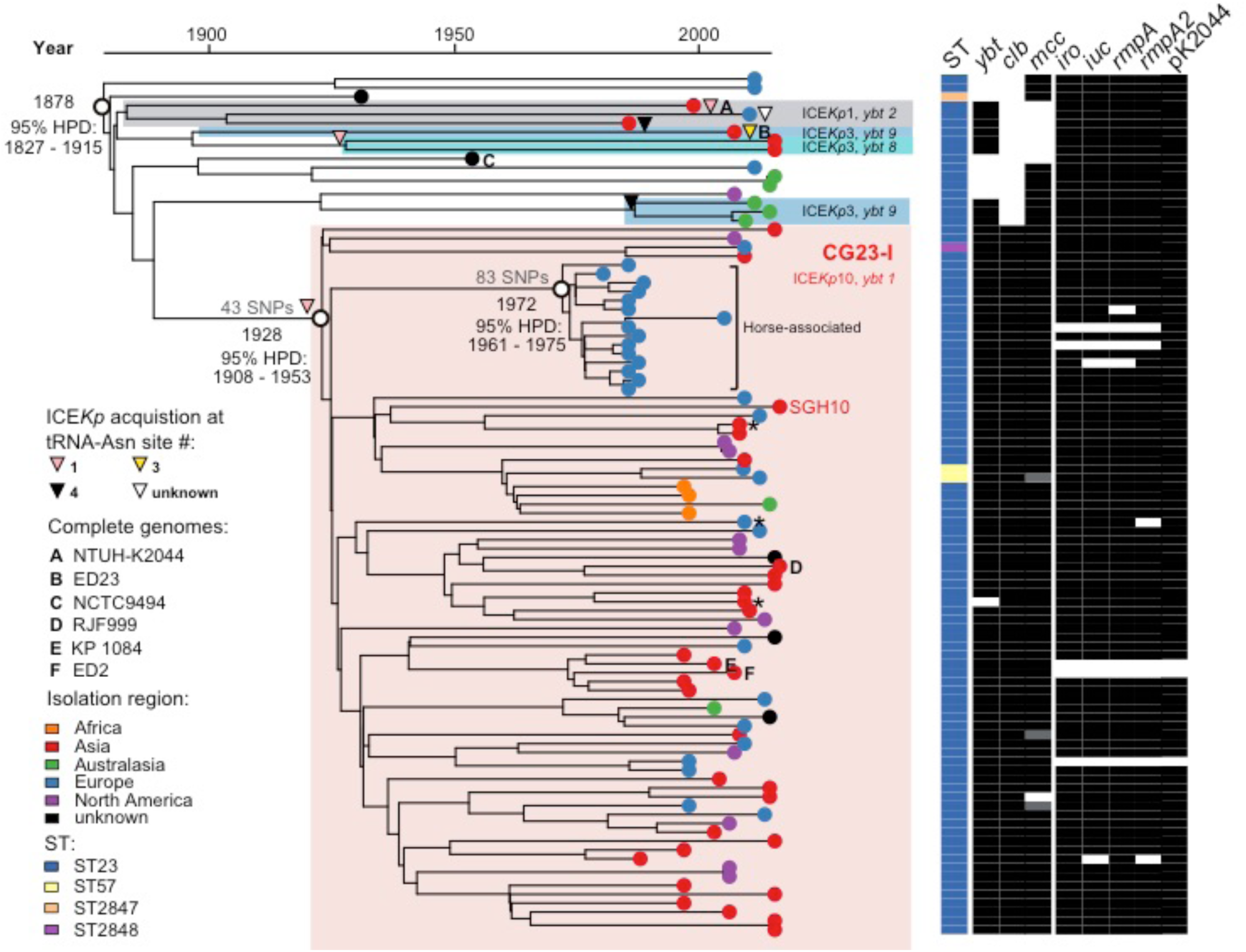
Phylogenetic relationships within CG23. Time-calibrated Bayesian phylogeny (left) showing the relationships between 97 CG23 *Kp*, their sequences types (STs), the presence of virulence loci (*ybt*, yersiniabactin;*clb*, colibactin; *mcc*, microcin E492; *iro*, salmochelin; *iuc*, aerobactin; *rmpA/rmpA2*, regulator of mucoid phenotype) and the virulence plasmid (pK2044-like, see **Fig. S6**).

We detected a strong temporal signal in the CG23 alignment (**Fig. 2A-C, Supplementary Methods**), sufficient to estimate substitution rates and divergence dates for the CG23 population. The best-fitting Bayesian analysis using BEAST (UCLD constant model, see Supplementary Methods) estimated the mean evolutionary rate to be 3.40×10^-7^ substitutions site^-1^ year^-1^ (95% HPD; 2.43×10^-7^ - 4.38×10^-7^). The most recent common ancestor (MRCA) for all CG23 was estimated at 1878 (95% HPD; 1827-1915) and the MRCA for CG23-I at 1928 (95% HPD; 1908-1953) (**Fig. 1, Fig. 2D-F**). Importantly, there was significant overlap in the Bayesian parameter estimate distributions from four alternative models, and with estimates derived from an alternative method known as least-squares LSD analysis (**Fig. 2D-F**, see **Supplementary Methods**). This provides confidence that CG23 emerged some time in the late 19^th^ century and the globally distributed CG23-I emerged in the 1920s. The MRCA for the equine subclade nested within CG23-I was estimated at 1972 (BEAST analysis; 95% HPD, 1961-1975).

**Figure 2:**
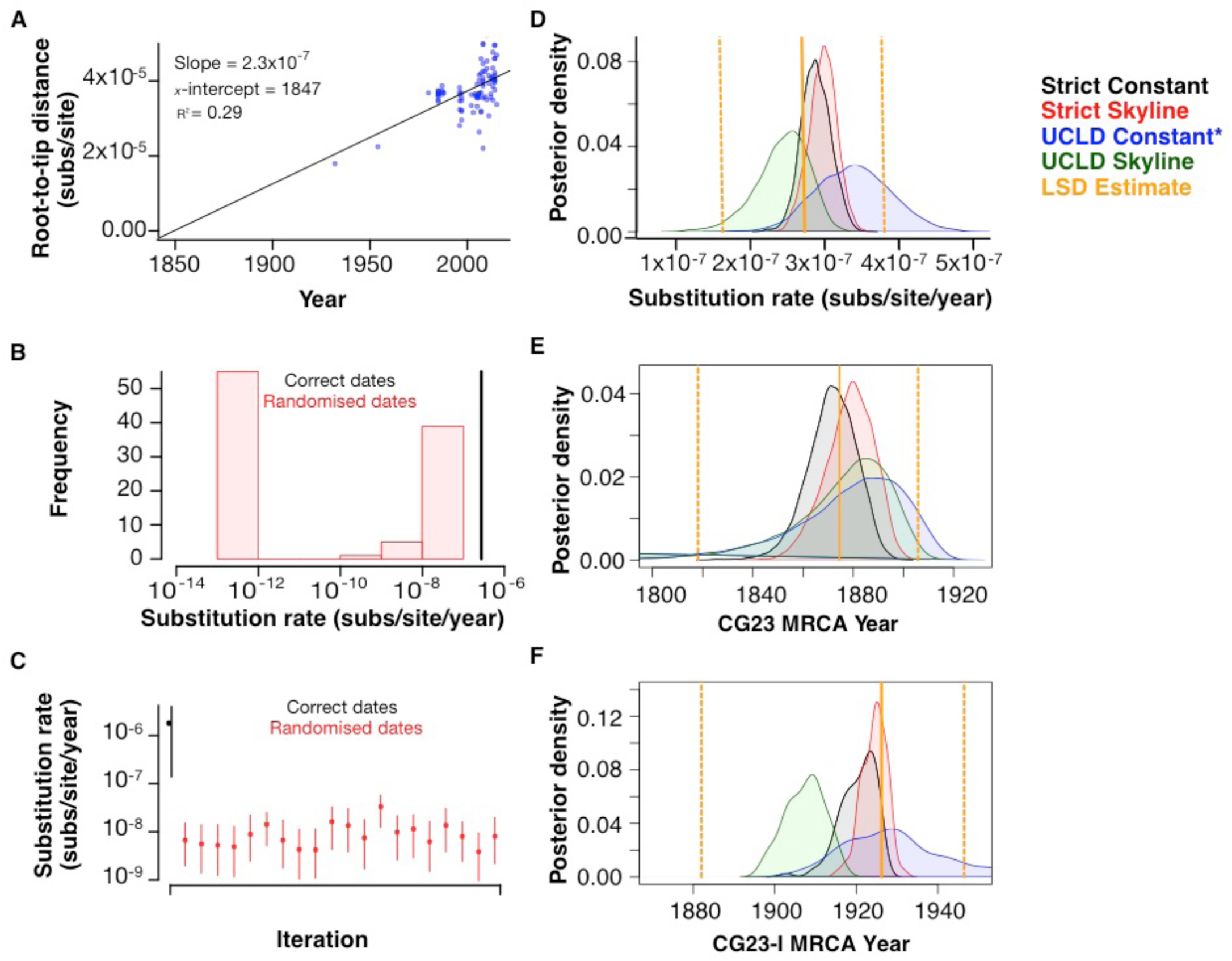
Evidence for temporal signal with agreement of dating methods and models. **A)** Correlation between root-to-tip distance (ML phylogeny) and year of isolate collection. **B)** Substitution rate estimates generated by the LSD least squares method with the correct isolate collection dates (black) and randomised dates (red, n = 100 tests). **C)** Posterior distributions for substitution rate estimates generated by the Bayesian method implemented in BEAST with the correct isolate collection dates (black) and randomised dates (red, n = 20 tests). **D, E and F)** Distributions of estimates for substitution rates, time to most recent common ancestor (TMRCA) for all CG23 and TMRCA for CG23-I, respectively. Distributions for each of four different Bayesian models are distinguished by colour as indicated, with the model used for the phylogeny in **Fig. 1, S4 and S6** marked with an asterisk. Estimates from the LSD method are shown in yellow +/− 95% confidence interval.

We hypothesised that the successful spread of CG23-I in the human population may have been associated with an increase in diversification rate and effective population size, compared to the rest of the CG23 population. To test this we investigated birth/death rates for nested subclades in the tree (the ratio of lineages that arise vs those that go extinct; a ratio >1 indicates population expansion, <1 indicates population decline, see **Supplementary Methods**). We first examined CG23-I, and compared the equine subclade to the rest of CG23-I (human isolates); this indicated population decline in the horse clade and expansion in the human CG23-I population (**Fig. S3A,C**). We then pruned the horse isolates from the tree, to consider only human isolates, and compared the CG23-I isolates to the rest of the CG23 population (i.e. non-CG23-I). This provided evidence for population expansion amongst human-associated CG23 generally, but with >5x greater expansion in CG23-I than the rest of the population (**Fig. S3B,D**).

### Genome evolution and gene content variation in the CG23 population

Comparison of seven completely assembled CG23 chromosomes (isolated between 1954 and 2015, including a newly completed genome for strain SGH10) revealed strong conservation of genome structure, interrupted only by the acquisition of ICEs or prophage, and two inversions (one mediated by an IS, one by phage, see **Fig. 3, Supplementary Results**). Pangenome analysis of the full set of 97 genomes identified 4170 core genes (present in ≥95% of genomes) and 5493 accessory genes (present in <10% of genomes and mostly associated with phage and plasmids, see **Fig. S4, Supplementary Results**). Nineteen coding sequences were uniquely present in CG23-I (>95% CG23-I genomes). Eighteen of these were part of the colibactin synthesis locus *clb* (detailed below) and the other was an IS*Kpnl* transposase inserted within an ethanolamine transporter gene, *eat* (KP1_1165 in NTUH-K2044). Most (95%) of the CG23 genomes harboured an intact CRISPR/Cas locus and we identified extensive spacer diversity in the corresponding CRISPR arrays, suggesting the system is active in defending against foreign DNA (see **Supplementary Results**).

**Figure 3:**
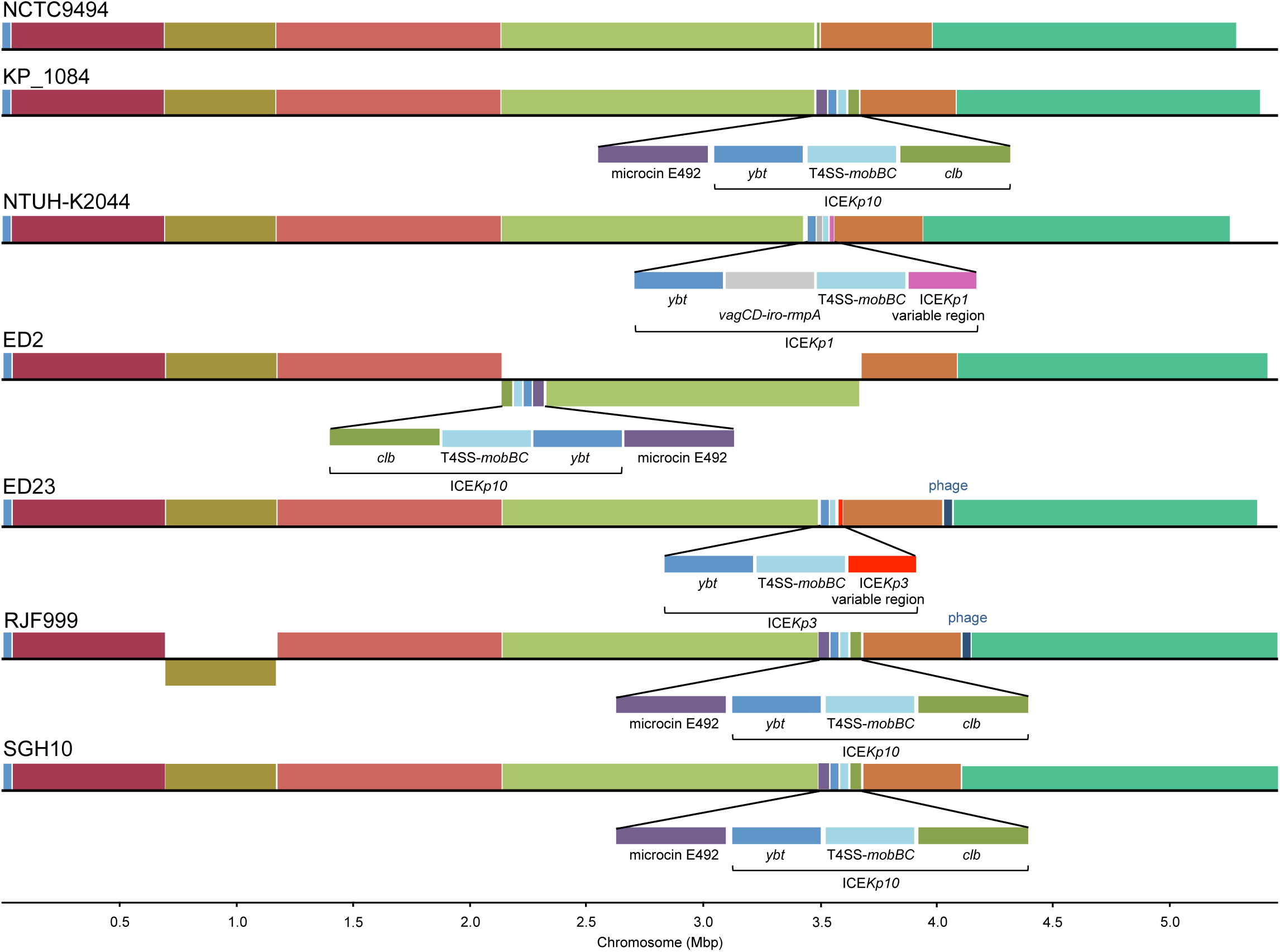
Chromosomal synteny and content comparisons between CG23 completed assemblies. Homologous regions common to the chromosomes for six completed CG23 chromosomes are represented as blocks. Chromosomal inversions relative to the oldest genome NCTC9494 are indicated by blocks that sit below the centre line for a corresponding genome, while those that sit above the line indicate the same orientation as the reference. Unique acquisitions that were not common to all genomes are annotated.

To investigate the ongoing evolution of virulence in CG23, we performed detailed analyses of variation in virulence gene content within the population. All CG23 genomes (including the hybrid strain CAS686) carried the KL1 locus associated with biosynthesis of the K1 capsule serotype. The K-locus of the 1954 isolate NCTC9494 harboured a 1065 bp insertion between the *wzi* and *wza* genes, which contained an IS*1O2*-like transposase; insertions in K-loci are common amongst historical isolates ^38^ and likely arise during long-term storage.

The *ybt* locus was present in 90 CG23 genomes (93%). Comparison of *ybt* lineages, ICE*Kp* structures and integration sites indicated at least six independent *ybt+* ICE*Kp* acquisitions (triangles in **Fig. 1**). CG23-I was characterised by the presence of ICE*Kp10* encoding *ybt* lineage 1 (*ybt*-1) and *clb* lineage 2A sequence variants, integrated at tRNA-Asn site 1, consistent with a single integration event prior to the expansion of the sublineage. One isolate from this group lacked the *ybt* genes (**Fig. 1**) but harboured the left and right ends of ICE*Kp10* (**Fig. S5**), consistent with this genome sharing the historical acquisition of ICE*Kp10* followed by subsequent loss of *ybt* and the ICE*Kp* mobilisation machinery.

At least five unrelated ICE*Kp* integration events were identified outside of CG23-I (**Fig. 1**), none of which included *clb*. Three separate introductions of ICE*Kp3* were detected: ED23 (from Taiwan) carried *ybt*-9 integrated at tRNA-Asn site 3, three isolates from Australia carried *ybt*-9 variant integrated at tRNA-Asn site 4, and two related Singapore isolates carried *ybt*-8 integrated at tRNA-Asn site 1. ICE*Kp1* encoding *ybt*-2 was identified in two strains from Taiwan (including the NTUH-K2044 reference strain) and one strain from France.These strains were monophyletic in the tree (grey shaded clade in **Fig. 1**) and share a MRCA close to the root, however the locations of ICE*Kp1* suggest they may have been acquired via distinct integration events in each strain since their divergence: at site 1 in NTUH-K2044 and site 4 in SB3926 (the integration site could not be resolved for the French isolate SB4446).

The virulence plasmid-associated loci *iuc*, *iro*, *rmpA* and *rmpA2* were present in the vast majority of CG23 genomes (see Fig. 1). Comparison of genome sequences to the reference sequence of virulence plasmid pK2044 (strain NTUH-K2044) confirmed that the plasmid backbone was present in 94 genomes, including all those carrying *iuc*, *iro*, *rmpA* and/or *rmpA2*, although some plasmids harboured deletions that affected these virulence loci (see **Fig. 1, Fig. S6A**; note it is unclear whether the three plasmid-negative isolates were actually lacking the plasmid *in vivo* or had lost it during laboratory culture). Other virulence loci detected in CG23 included microcin E492 (three deletion variants detected, see **Fig. S7, Supplementary Results**) and allantoinase, both of which were found in the chromosome.

### SGH10 as a novel reference strain for hypervirulent CG23

Our data on CG23 population structure and virulence loci show that none of the currently available finished genome sequences represent a ‘typical’ genome. NTUH-K2044 (liver abscess), ED23 (blood) and NCTC 9494 (sputum) are not part of the predominant CG23-I sublineage and do not harbour the colibactin locus (labelled A-C in **Fig. 1**). Strains 1084 (liver abscess) and ED2 (blood) belong to CG23-I but lack the virulence plasmid, and RJF999 is a blood isolate of undetermined virulence. We therefore propose strain SGH10, which belongs to the predominant CG23-I sublineage and has all virulence loci intact and no atypical accessory genes (see **Fig. 1** and **S4**), as a reference strain for experimental and genomic studies of hypervirulent CG23 *K. pneumoniae* associated with human liver abscess.

SGH10 was isolated from a liver abscess patient in Singapore in 2014, and we have previously demonstrated it to be hypermucoid and highly resistant to human serum ^19^. Here we confirmed the virulence potential of SGH10 via oral infection of C57BL/6 mice, which resulted in translocation to the liver, lungs and spleen 48 hours post-inoculation (see **Methods** and **Fig. 4A**). We performed additional long and short read sequencing of the SGH10 genome (see Methods), yielding complete circular sequences for the chromosome (5,485,114 bp, 57.43% GC content) (**Fig. 4B**) and virulence plasmid (pSGH10; 231,583 bp, 50.15% GC) (**Fig. 4C**). The annotated genome sequence was deposited in GenBank under accessions CP025080 (chromosome) and CP025081 (virulence plasmid), and the strain submitted to the National Collection of Type Cultures in the United Kingdom (NCTC number 14052).

**Figure 4:**
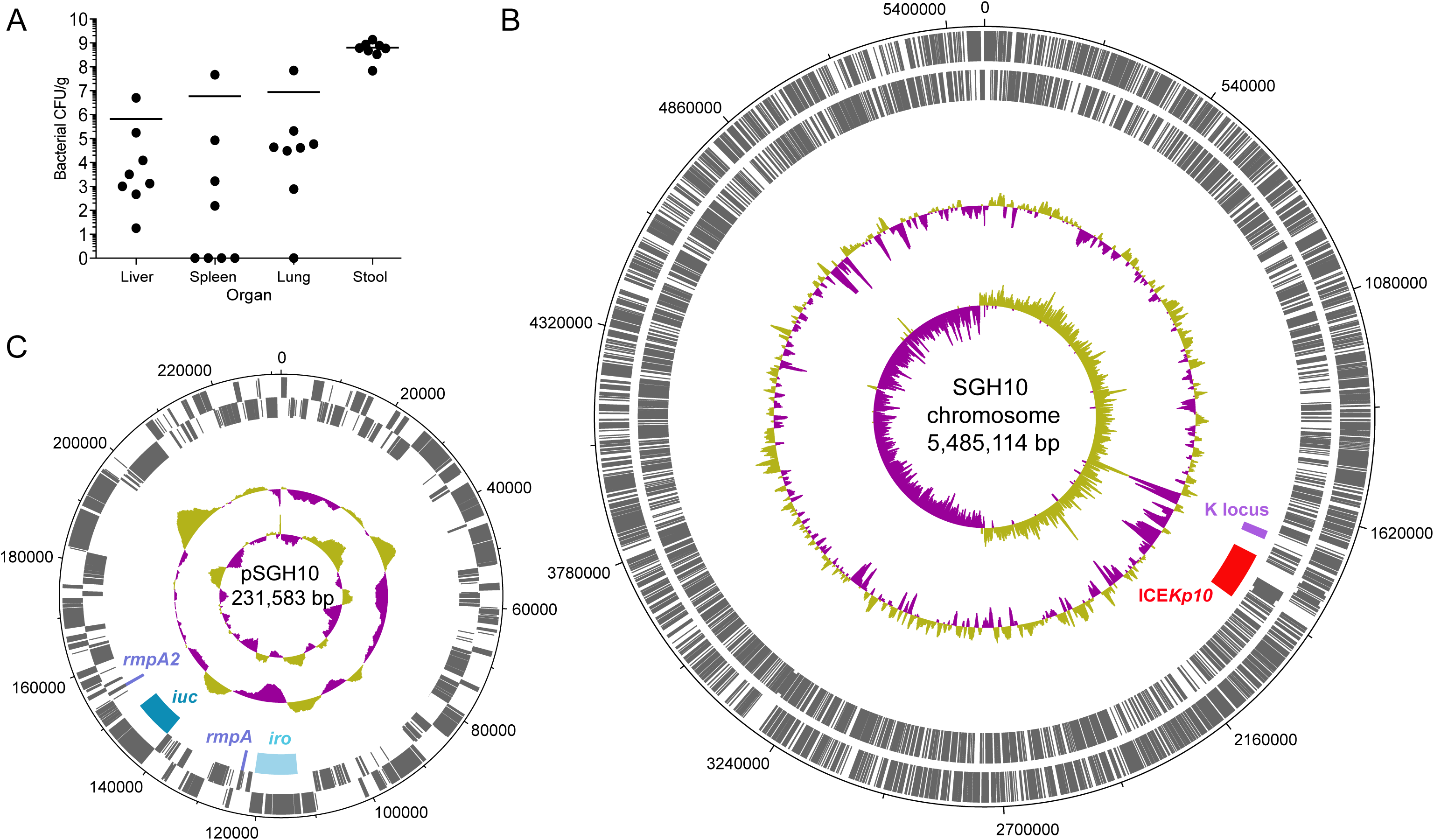
Virulence and genomic properties of proposed CG23-I strain SGH10. **A)** Bacterial burden (CFU/g) in organs and stool of n=8 mice 48 hours following oral infection. **B)** SGH10 chromosome and **C)** plasmid. Tracks shown are (from inner to outer): GC skew (G-C/G+C), G+C content, key capsule and virulence features as labelled, coding sequences on reverse strand and coding sequences on forward strand.

### Antimicrobial resistance determinants in CG23

All CG23 genomes carried the intrinsic beta-lactamase gene *bla*_SHV_ (which confers ampicillin resistance) and the *oqxAB* genes (which confer reduced susceptibility to quinolones).

However no fluoroquinolone resistance-associated mutations were identified in *gyrA* or *parC,* and acquired AMR determinants were rare. The equine clade and three other unrelated CG23-I isolated from humans (CAS813, BG130 and BG141; * in **Fig. 1**) carried multiple AMR genes. Variants of an IncFII plasmid, pSB4816, conferred multidrug resistance in the equine clade (see **Fig. S8, Supplementary Results**). The multidrug-resistant strains isolated from humans each harboured unique sets of AMR genes conferring resistance to multiple classes of drugs, as previously reported ^27,28^ (see **Table S1**). While precise locations of the AMR genes were not resolvable in the draft genome assemblies, all three were found to harbour large conjugative plasmids that are frequently associated with multidrug resistance in Enterobacteriaceae: IncA/C_2_ plasmid sequence type (PST) 3 in CAS813 (Denmark, 2008) and BG130 (Vietnam, 2008); IncN PST6 in BG141 (Madagascar, 2007).

## Discussion

The earlier comparative genomic studies of up to 27 CG23 ^27,28^ included only four genomes outside of CG23-I, which somewhat obscured the diversity and population structure of the clonal group, and provided insufficient temporal signal to reconstruct evolutionary dynamics. Here, the expanded genome collection revealed that the majority of human clinical isolates of CG23 belong to a clonally expanded sublineage, CG23-I, which emerged in the early 20^th^ century (**Fig. 1**). This predates the identification of serotype K1 *Kp* as a significant cause of liver abscess by over 50 years ^4,39^, suggesting it has been circulating undetected for many decades. We also found evidence that the entire population of CG23 associated with human infections has been expanding since its emergence, but that the CG23-I sublineage has undergone particularly accelerated population expansion and global dissemination (**Fig. S3**).

Our genomic analyses suggest some potential mechanisms for the success of the CG23-I sublineage in the human population. The colibactin synthesis locus *clb* (also referred to as *pks*), encoded downstream of the *ybt* locus in ICE*Kp10*, is its most notable feature. Colibactin is a hybrid nonribosomal peptide-polyketide that has a genotoxic effect on host cells by crosslinking DNA and inducing double strand DNA breaks ^31,40,41^. It was first discovered in *E. coli* ^40^ but has since been reported in 3.5-4% of *K. pneumoniae* ^26,42^, in which it has also been shown to induce DNA double strand breaks in HeLa cells ^42^. Genotoxicity of colibactin in ST23 *Kp* has been demonstrated for strain 1084 (which belongs to CG23-I), both *in vitro* in mouse liver cells and *in vivo* in liver parenchymal cells of orally infected BALB/c mice ^31^. In these experiments, the genotoxic effect of the ST23 *Kp* strain was clearly attributed to colibactin production, using isogenic Δ*clbA* and complemented mutants ^31^. Subsequent infection experiments using the same strains revealed that loss of colibactin caused a reduction in dissemination to the blood, liver, spleen and brain ^43^. Crucially, this work also indicated that the Δ*clbA* mutant was attenuated in its ability to colonise the intestinal mucosa, which is considered a critical prerequisite for invasive disease ^43^. Similarly, colibactin has been shown to promote gut colonisation and to be essential for disseminated infection by *E. coli* ^41,44^ Therefore, the acquisition of the *clb* locus likely promotes both gut colonisation and mucosal invasion of CG23-I *Kp,* leading to enhanced transmissibility and increased virulence, which may explain the dissemination of the lineage and its dominance amongst hypervirulent infections globally. Concerningly, colibactin also increases the likelihood of serious complications such as metastatic spread to the brain ^43^ and potentially tumorigenesis ^45-48^. Colibactin-positive *Kp* is particularly common in Taiwan, where it has been reported in 17-25% of non-abscess infections and significantly associated with K1 strains (which are most likely ST23) ^45,49^. In this setting pyogenic liver abscess, particularly caused by *K. pneumoniae,* has been shown to be a significant risk factor for colorectal cancer ^45,50^ and liver cancer ^51^. K1 and/or ST23 *Kp* gut colonisation rates up to 10% have been reported across East Asia ^52,53^. The prevalence of colibactin among these strains has not yet been investigated in most of these countries, however *Kp* liver abscess has been associated with colorectal cancer across East Asia^54,55^, and based on our genome data we predict the majority of the strains in question will belong to the predominant *clb*+ CG23-I sublineage.

The only other gene content changes characteristic of CG23-I were loss-of-function mutations in the fimbrial protein KpcC (via nonsense mutation) and the ethanolamine transporter Eat (via IS insertion). Whether these differences in surface structures or metabolic pathways reflect adaptive selection, or simply loss of functions that are not advantageous to the clone, remains a topic for future research. *Kp* has two operons involved in ethanolamine metabolism: *eutRKLCBAHGJENMDTQP* and *eat-eutBC*. The former is typically associated with facultative anaerobes that live in the gut and/or mouth, and the latter with obligate aerobes that may be found in the environment ^56,57^; hence the loss of Eat may reflect adaptation to the host-associated, anaerobic niche. A recent metabolic study of *Kp* growth under aerobic conditions included two ST23 strains NTUH-K2044 (intact *eat)* and SB4385 (CG23-I, disrupted *eat*). Both grew on ethanolamine with no apparent differences in efficiency ^58^, indicating no detrimental effect of the loss of Eat in CG23-I strains during aerobic growth, hence the available evidence indicates this change is unlikely to be of particular functional significance to the lineage.

The K1 capsule synthesis locus KL1, microcin ICE, and virulence plasmid encoding aerobactin, salmochelin and *rmpA* (**Fig. 1, S6**) were present in most strains, suggesting they were in the ancestor of CG23 and have been selectively maintained. In contrast, distinct variants of ICE*Kp* have been acquired on multiple separate occasions after divergence from the CG23 ancestor. Notably only one such acquisition – that of *clb*-positive ICE*Kp10* in CG23-I – was associated with subsequent clonal expansion, suggesting positive selection for colibactin production. The available data suggest that no particular virulence factor of CG23 is necessary or sufficient for invasive disease. Three liver abscess strains lacked ICE*Kp*, and it was recently reported that the 1932 Murray Collection isolate lacking ICE*Kp* is highly virulent in the *Galleria mellonella* infection model ^37^. Both asymptomatic carriage isolates in our study lacked ICEKp, however the significance of this is not yet clear. Asymptomatic gastrointestinal carriage of K1 CG23 has been reported previously, with up to 10% prevalence in Asian populations ^52,53^, but whether these carry ICE*Kp10* or belong to CG23-I is unknown. CG23-I strain 1084, which lacks the virulence plasmid but carries *ybt* and *clb* in ICE*Kp10*, was isolated from human liver abscess and is reportedly highly invasive in murine models of pneumonia and liver abscess ^31,59^.

The emergence of AMR CG23 is a significant potential health threat. Treatment of pyogenic liver abscess relies on drainage of the abscess and effective antimicrobial therapy^1^. AMR has been occasionally reported in human isolates of CG23 ^32-34,60^, and our genomic analysis confirmed that acquisition of AMR plasmids was rare, with distinct plasmids identified in just three sporadic strains from human infections ^27,28^ and no evidence of long-term plasmid maintenance or transmission amongst human isolates (**Fig. 1**). This is in contrast to the frequency, diversity and apparent stability of AMR plasmids found in other *Kp* clones such as CG258 or CG15 ^61-54^, which also show evidence of frequent ICE*Kp* integration ^26^. Overall, our population genomics data suggest there may be barriers to plasmid acquisition and maintenance in CG23 that have so far protected us against the emergence of widespread AMR in this hypervirulent clone. These may include the hyper-expression of K1 capsule, which may provide a physical barrier against transformation and conjugation and also CRISPR/Cas systems, which defend against foreign DNA that does manage to penetrate the cell. The significant association between upregulated capsule production and reduced uptake of DNA has previously been documented in *Streptococcus pneumoniae* ^65^. Notably, these mechanisms could also explain the lack of homologous recombination in CG23, which is common in other clonal groups ^66,67^. However, the occasional acquisition of plasmids, including AMR plasmids, in CG23 shows that these barriers are not complete and the maintenance of an IncFII AMR plasmid in the equine clade over 20 years (**Fig. S6, S8**) shows that long-term stability of AMR plasmids is possible in CG23. Hence our data indicate we must anticipate and carefully monitor for the emergence of stable AMR in CG23.

Taken together our findings have important implications for future study of hypervirulent *Kp*. Firstly, we suggest that epidemiological studies of *Kp* liver abscess infections should investigate and report not only chromosomal MLST and virulence plasmid marker genes such as *rmpA*, but also the *ybt* and *clb* loci, which can be used as markers for the CG23-I sublineage. These loci can be rapidly identified in genome data using our *Klebsiella* genotyping tool, *Kleborate* (https://github.com/katholt/Kleborate/)or can be identified by conventional PCR (primers in ref ^19^). Secondly, our observations regarding the population structure and virulence loci show that the two strains that have been used as reference genomes and/or experimental models for hypervirulent ST23 associated with liver abscess are rather atypical: NTUH-K2044 ^29,31^ does not belong to the dominant sublineage CG23-I and lacks colibactin, while 1084 ^31,68^ lacks the virulence plasmid. We therefore suggest that future studies aiming to explore the virulence, transmissibility or evolution of CG23 *Kp* should consider genome-based profiling prior to the selection of clinical isolates for experimental work. In addition, we present the human liver abscess isolate SGH10 as an open access reference strain for future studies into hypervirulent CG23 *Kp* associated with liver abscess (strain available under NCTC 14052; complete genome available under GenBank accessions CP025080, CP025081). SGH10 belongs to the globally expanded GC23-I sublineage, carries intact copies of all virulence loci, is hypermucoid and serum-resistant, and is capable of causing disseminated infection in a murine model. Finally, we advocate that researchers, clinicians and public health authorities co-ordinate efforts to improve the surveillance of hypervirulent *Kp* and facilitate rapid identification of emerging threats, such as convergence of hypervirulence and AMR, that may otherwise remain undetected for decades.

## Methods

### Genome collection

We identified 83 CG23 genome sequences from our curated collection (six finished and 77 draft genomes, see **Table S1** ^26,38^). These included 43 genomes from our own previous studies ^19,21,27,69^ and 40 publicly available genomes including the 27 analysed by Struve *et al* ^28,29,37,68,70-74^. We also sequenced 15 novel genomes that were identified as ST23 by MLST (02A029, 12A041 and 16A151 ^14^) or identified as K1 by classical serotyping and then confirmed as ST23 by genome sequencing (12 strains isolated between 1980 and 1987 from samples from horses; see **Table S1**). DNA was extracted by the phenol-chloroform method, libraries prepared using Nextera technology and paired end reads of either 100 bp (Illumina HiSeq 2000) or 300 bp (Illumina MiSeq) were generated. Accession numbers for all 98 genomes included in this work are listed in **Table S1** (note though that the hybrid strain CAS686 was excluded from most analyses, as explained below).

### Virulence and genomic characterisation of novel CG23-I reference strain SGH10

We previously sequenced SGH10 (liver abscess isolate, Singapore 2015) via Illumina MiSeq^19^. Here we generated additional short read (Illumina MiniSeq) and long read (Oxford Nanopore) sequences (see **Supplementary Methods**) and constructed a high quality finished genome sequence using our hybrid assembler Unicycler^75^. Virulence of SGH10 was assessed using a murine oral infection model as previously described^19^ with slight modification (see **Supplementary Methods**).

### Genotyping, integrative and conjugative element (ICE) and K-locus typing

Where available (n = 89), short read sequence data were *de novo* assembled using Unicycler v0.3.0b ^75^ with SPAdes v3.8.1 ^76^ and annotated with Prokka v1.11 ^77^. SRST2 ^78^ was used to (i) determine chromosomal, yersiniabactin and colibactin multi-locus sequence types (STs) ^18,26^ and (ii) screen for virulence genes included in the BIGSdb ^27^, acquired AMR genes in the ARG-Annot database ^79^ and plasmid replicon genes in the PlasmidFinder database ^80^. Nine of the publicly-available genomes were available only as pre-assembled sequences. For these we used *Kleborate*(https://github.com/katholt/Kleborate/)to determine MLST, virulence and AMR information, and identified plasmid replicons separately by BLASTn search of the assemblies (coverage ≥90%, identity ≥90%; databases as above). Genomes with novel alleles were submitted to the *Kp* BIGSdb (http://bigsdb.pasteur.fr/klebsiella/klebsiella.html/)for assignment of novel allele and ST numbers.

ICE*Kp* structures were predicted for all genome assemblies using *Kleborate* ^26^ and confirmed by manual inspection using the Artemis genome browser ^81^. ICE*Kp* integration sites were determined (from the four possible chromosomal tRNA-Asn sites) using the Bandage assembly graph viewer ^82^ and BLASTn searches. Microcin E492 ICE sequences were extracted based on their flanking direct repeats and BLASTn searches were used to confirm their integration at tRNA-Asn site 2. K-loci were typed from the assemblies using *Kaptive* ^38^.

### Phylogenetic inference and molecular clock analyses

Sequence reads were mapped to the NTUH-K2044 reference chromosome (accession: AP006725) and single nucleotide variants were called using the RedDog v10b pipeline (https://github.com/katholt/reddog/)as described in **Supplementary Methods**. The previous study by Struve *et al* ^28^ showed that isolate CAS686 (ST260) is a hybrid recombinant strain, for which only half of the genome is closely related to CG23. Our mapping data confirmed this, and we further compared the CAS686 genome to our non-CG23 *Kp* collection ^26,38^ to identify the potential donor lineage (see **Supplementary Results** and **Fig. S1** for further details.) We therefore excluded CAS686 from our main phylogenetic and comparative analyses of CG23, which focus on the remaining 97 CG23 genomes, amongst which 6,838 variant sites were identified. The alignment of these sites was screened for further recombination using *Gubbins* ^83^, which did not detect any plausible recombination events. We used the complete 6,838 site alignment to infer a ML phylogeny using RAxML v8.1.23 ^84^ with the GTR+ nucleotide substitution model. The reported ML tree is that with the highest likelihood out of five independent runs. To assess branch support, we conducted 100 non-parametric bootstrap replicates using RAxML. The ML tree and alignment were used as input to FastML ^85^ for ancestral state reconstruction along the tree.

We used TempEst ^86^ to investigate the relationship between root-to-tip distances in the ML tree and year of isolation. We then used two different methods to infer the evolutionary rate and timescale (see **Supplementary Methods** for details). The first method consists of least-squares dating, implemented in LSD v0.3 ^87^, using as input the ML tree and year of isolation data. We then analysed the data using the Bayesian framework in BEAST v1.8 ^88^, using two clock models (strict and uncorrelated log normal (UCLD)) and two demographic models (constant population size and Bayesian skyline), as detailed in **Supplementary Methods**. UCLD clock with constant population size was selected as the best model. We additionally used BEAST to test whether diversification rates differed among lineages (see **Supplementary Methods**).

### Structural rearrangements, pan-genome analyses and CRISPR array variation

The chromosomal organisation of the six publicly available completely assembled ST23 genomes (NTUH-K2044, 1084, NCTC9494, ED2, ED23 and RJF999; accessions in **Table S1**) and our newly completed SGH10 genome were subjected to multiple alignment and visualisation using Mauve ^89^. All 97 genome assemblies were screened for the presence of phage using PHAST ^90^. The pan-genome was investigated using Roary^91^ with the Prokka-annotated genomes as the input and ≥95% amino acid identity for protein clustering. A single representative sequence for each gene cluster was compared to the putative phage sequences and the novel pSB4816 sequence (see **Supplementary Methods**) using BLASTn, in order to identify the components of the pan-genome belonging to phage and pSB4816, respectively. CRISPR arrays were identified using the CRISPR Recognition Tool ^92^ and *cas* operon genes were identified by tBLASTx (id ≥ 90%, coverage ≥ 28% compared to those annotated in the NTUH-K2044 genome). CRISPR spacer sequences were extracted and clustered at 100% nucleotide identity using CD-HIT-EST ^93^.

### Plasmid analyses

To assess the conservation of the pK2044 virulence plasmid, sequence reads were mapped to the pK2044 reference sequence (accession: AP006726.1) using RedDog, and each annotated gene was counted as present in a given isolate if the reads covered ≥90% of the gene length at read depth ≥5. Putative AMR plasmids were investigated using the Bandage assembly graph viewer ^82^ as detailed in **Supplementary Methods** for details. We were able to extract a novel plasmid sequence, pSB4816, from the assembly graph of equine isolate SB4816, which was resolved into two contigs. Conservation of this plasmid was assessed by mapping each read set to the concatenated pSB4816 sequence as above. Unfortunately we were not able to extract the complete sequences of any other AMR plasmids due to complexities in the assembly graphs, but we were able to identify the putative plasmid replicon types and associated plasmid STs (see **Supplementary Methods**).

### Data availability

Accession numbers for all genome data included in this work are summarised in **Table S1**. Illumina sequence reads for the 15 newly sequenced equine isolates have been deposited in the NCBI sequence read archive under accession PRJNA391004. The finished genome sequence for SGH10 (BioSample; SAMN06112188) was deposited in Genbank (accessions CP025080, CP025081) and the strain was deposited in the UK National Collection of Type Cultures (NCTC 14052); novel Illumina and Oxford Nanopore sequence reads are available in the sequence read archive under accessions SRR6266394, SRR6266393, SRR6266392 and SRR6307304. The sequence for plasmid pSB4816 was deposited in GenBank (accessions: MF363048, MF966381). The Bayesian phylogenetic tree, annotated with strain information,is available for download and interactive visualisation at https://microreact.org/project/r1_dSC9w-/.

## Acknowledgments

We thank Jonathan Cotrupi for microbiological analysis of horse strains from France and Ulises Garza-Ramos and Alexis Criscuolo for help in genomic analyses. We also thank the team of the curators of the Institut Pasteur MLST system (Paris, France) for importing novel alleles, profiles and/or isolates at http://bigsdb.pasteur.fr./ This work was supported by the NHMRC of Australia (fellowship #1061409 to KEH).

## References

1. Shon, A. S., Bajwa, R. P. S. & Russo, T. A. Hypervirulent (hypermucoviscous) *Klebsiella pneumoniae:* a new and dangerous breed. Virulence 4, 107–18 (2013).

2. Siu, L. K., Yeh, K., Lin, J., Fung, C. & Chang, F. *Klebsiella pneumoniae* liver abscess: a new invasive syndrome. Lancet Infect Dis 12, 881–887 (2012).

3. Lee, K. H., Hui, K. P., Tan, W. C. & Lim, T. K. *Klebsiella* bacteraemia: a report of 101 cases from National University Hospital, Singapore. J Hosp Infect 27, 299–305 (1994).

4. Cheng, D., Liu, Y., Yen, M., Liu, C. & Wang, R. Septic Metastatic Lesions of Pyogenic Liver Abscess Their Association With *Klebsiella pneumoniae* Bacteremia in Diabetic Patients. Arch Intern Med 151, 1557–1559 (1991).

5. Qian, Y. et al. A retrospective study of pyogenic liver abscess focusing on *Klebsiella pneumoniae* as a primary pathogen in China from 1994 to 2015. Sci Rep 6, 38587 (2016).

6. Chung, D. R. et al. Emerging invasive liver abscess caused by K1 serotype *Klebsiella pneumoniae* in Korea. J Infect 54, 578–583 (2007).

7. Ko, W. et al. Community-Acquired *Klebsiella pneumoniae* Bacteremia: Global Differences in Clinical Patterns. Emerg Infect Dis 8, 160–166 (2002).

8. Dulku, G. & Tibballs, J. Cryptogenic invasive *Klebsiella pneumoniae* liver abscess syndrome (CIKPLA) in Western Australia? AMJ 7, 436–440 (2014).

9. Holmas, K., Fostervold, A., Stahlhut, S. G., Struve, C. & Holter, J. C. Emerging K1 serotype *Klebsiella pneumoniae* primary liver abscess: three cases presenting to a single university hospital in Norway. Clin Case Rep 2, 122–127 (2014).

10. Decre, D. et al. Emerging Severe and Fatal Infections Due to *Klebsiella pneumoniae* in Two University Hospitals in France. J Clin Microbiol 49, 3012–3014 (2011).

11. Frazee, B. W., Hansen, S. & Lambert, L. Invasive Infection With Hypermucoviscous *Klebsiella pneumoniae:* Multiple Cases Presenting to a Single Emergency Department in the United States. Ann Emerg Med 53, 639–642 (2009).

12. Pastagia, M. & Arumugam, V. *Klebsiella pneumoniae* liver abscesses in a public hospital in Queens, New York. Travel Med Infect Dis 6, 228–233 (2008).

13. Coutinho, R. L. et al. Community-acquired invasive liver abscess syndrome caused by a K1 serotype *Klebsiella pneumoniae* isolate in Brazil: a case report of hypervirulent ST23. Mem Inst Oswaldo Cruz 109, 970–971 (2014).

14. Brisse, S. et al. Virulent clones of *Klebsiella pneumoniae:* Identification and evolutionary scenario based on genomic and phenotypic characterization. PLoS One 4, (2009).

15. Luo, Y., Wang, Y., Ye, L. & Yang, J. Molecular epidemiology and virulence factors of pyogenic liver abscess causing *Klebsiella pneumoniae* in China. Clin Microbiol Infect 20, 0818–0824 (2014).

16. Liao, C. H., Huang, Y. T., Chang, C. Y., Hsu, H. S. & Hsueh, P. R. Capsular serotypes and multilocus sequence types of bacteremic *Klebsiella pneumoniae* isolates associated with different types of infections. Eur J ClinMicrobiol Infect Dis 33, 365–369 (2014).

17. Turton, J. F. et al. Genetically similar isolates of *Klebsiella pneumoniae* serotype K1 causing liver abscesses in three continents. J Med Microbiol 56, 593–597 (2007).

18. Diancourt, L., Passet, V., Verhoef, J., Grimont, P. A. D. & Brisse, S. Multilocus Sequence Typing of *Klebsiella pneumoniae* Nosocomial Isolates. 43, 4178–4182 (2005).

19. Lee, I. R. et al. Differential host susceptibility and bacterial virulence factors driving *Klebsiella* liver abscess in an ethnically diverse population. Sci. Rep. 6, 29316 (2016).

20. Qu, T. et al. Clinical and microbiological characteristics of *Klebsiella pneumoniae* liver abscess in East China. BMC Infect Dis 15, 1–8 (2015).

21. Holt, K. E. et al. Genomic analysis of diversity, population structure, virulence, and antimicrobial resistance in *Klebsiella pneumoniae,* an urgent threat to public health. Proc Natl Acad Sci USA 112, E3574–81 (2015).

22. Nassif, X., Fournier, J., Arondel, J. & Sansonetti, P. J. Mucoid Phenotype of *Klebsiella pneumoniae* Is a Plasmid-Encoded Virulence Factor. Infect Immun 57, 546–552 (1989).

23. Bachman, M. A. et al. *Klebsiella pneumoniae* yersiniabactin promotes respiratory tract infection through evasion of lipocalin 2. Infect. Immun. 79, 3309–3316 (2011).

24. Holden, V. I., Breen, P., Houle, S., Dozois, C. M. & Bachman, M. A. *Klebsiella pneumoniae* Siderophores Induce Inflammation, Bacterial Dissemination, and HIF-1a Stablization during Pneumonia. MBio 7, e01397-16-10 (2016).

25. Hsieh, P., Lin, T., Lee, C., Tsai, S. & Wang, J. Serum-Induced Iron-Acquisition Systems and TonB Contribute to Virulence in *Klebsiella pneumoniae* Causing Primary Pyogenic Liver Abscess. J Infect Dis 197, 1717–1727 (2008).

26. Lam, M. M. C. et al. Frequent emergence of pathogenic lineages of *Klebsiella pneumoniae* via mobilisation of yersiniabactin and colibactin. bioXriv (2017). doi:https://doi.org/10.1101/098178

27. Bialek-davenet, S. et al. Genomic Definition of Hypervirulent and Multidrug-Resistant *Klebsiella pneumoniae* Clonal Groups. Emerg. Infect. Dis. 20, 1812–1820 (2014).

28. Struve, C. et al. Mapping the evolution of hypervirulent *Klebsiella pneumoniae*. MBio 6, 1–12 (2015).

29. Wu, K. M. et al. Genome sequencing and comparative analysis of *Klebsiella pneumoniae* NTUH-K2044, a strain causing liver abscess and meningitis. J. Bacteriol. 191, 4492–4501 (2009).

30. Lin, T. L., Lee, C. Z., Hsieh, P. F., Tsai, S. F. & Wang, J. T. Characterization of integrative and conjugative element ICEKp1-associated genomic heterogeneity in a *Klebsiella pneumoniae* strain isolated from a primary liver abscess. J. Bacteriol. 190, 515–526 (2008).

31. Lai, Y. C. et al. Genotoxic Klebsiella pneumoniae in Taiwan. PLoS One 9, (2014).

32. Cheong, H. S. et al. Emergence of serotype K1 *Klebsiella pneumoniae* ST23 strains co-producing the DHA-1 and an extended-spectrum beta-lactamase in Korea. Antimicrob. Resist. Infect. Control 2–5 (2016). doi: 10.1186/s13756-016-0151-2

33. Shin, J. & Ko, K. S. Single origin of three plasmids bearing bla CTX-M-15 from different *Klebsiella pneumoniae* clones. J Antimicrob Chemother 69, 969–972 (2014).

34. Liu, Y. et al. Capsular Polysaccharide Types and Virulence-Related Isolates in a Chinese University Hospital. Microb. Drug. Resist. (2017). doi: 10.1089/mdr.2016.0222

35. Wu, C., Huang, Y., Fung, C. & Peng, H. Regulation of the *Klebsiella pneumoniae* Kpc fimbriae by the site-specific recombinase KpcI. Microbiology 156, 1983–1992 (2010).

36. Samper, J. C. & Tibary, A. Disease transmission in horses. Theriogenology 66, 551–559 (2006).

37. Wand, M. E. et al. Characterization of pre-antibiotic era *Klebsiella pneumoniae* isolates with respect to antibiotic/disinfectant susceptibility and virulence in *Galleria mellonella*. Antimicrob. Agents Chemother. 59, 3966–3972 (2015).

38. Wyres, K. L. et al. Identification of *Klebsiella* capsule synthesis loci from whole genome data. Microb. Genomics (2016). doi:10.1099/mgen.0.000102

39. Liu, Y., Cheng, D. & Lin, C. *Klebsiella pneumoniae* liver abscess associated with septic endophthalmitis. Arch Intern Med 146, 1913–1916 (1986).

40. Vizcaino, M. I. & Crawford, J. M. The colibactin warhead crosslinks DNA. Nat Chem. 7, 411–417 (2015).

41. Nougayrède, J. P. et al. *Escherichia coli* Induces DNA Double-Strand Breaks in Eukaryotic Cells. Science (80-.). 313, 848–851 (2006).

42. Putze, J. et al. Genetic structure and distribution of the colibactin genomic island among members of the family Enterobacteriaceae. Infect. Immun. 77, 4696–4703 (2009).

43. Lu, M. et al. Colibactin Contributes to the Hypervirulence of*pks+* K1 CC23 *Klebsiella pneumoniae* in Mouse Meningitis Infections. Front. Cell. Infect. Microbiol. 7, (2017).

44. Mccarthy, A. J. et al. The Genotoxin Colibactin Is a Determinant of Virulence in *Escherichia coli* K1 Experimental Neonatal Systemic Infection. Infect Immun 83, 3704–3711 (2015).

45. Huang, W. et al. Higher rate of colorectal cancer among patients with pyogenic liver abscess with *Klebsiella pneumoniae* than those without: an 11-year follow-up study. Color. Dis. Off. J. Assoc. Coloproctology Gt. Britain Irel. 14, e794–801 (2012).

46. Raisch, J., Rolhion, N., Dubois, A., Darfeuille-Michaud, A. & Bringer, M. A. Intracellular colon cancer-associated *Escherichia coli* promote protumoral activities of human macrophages by inducing sustained COX-2 expression. Lab Invest 95, 296–307 (2015).

47. Cougnoux, A. et al. Bacterial genotoxin colibactin promotes colon tumour growth by inducing a senescence-associated secretory phenotype. Gut 63, 1837–1838 (2014).

48. Dalmasso, G., Cougnoux, A., Delmas, J., Darfeuille-Michaud, A. & Bonnet, R. The bacterial genotoxin colibactin promotes colon tumor growth by modifying the tumor microenvironment. Gut Microbes 5, 675–680 (2014).

49. Chen, Y. et al. Prevalence and characteristics of pks genotoxin gene cluster-positive clinical *Klebsiella pneumoniae* isolates in Taiwan. Sci Rep 7, (2017).

50. Kao, W. et al. Alimentary Pharmacology and Therapeutics Cancer risk in patients with pyogenic liver abscess: a nationwide cohort study. Aliment Pharmacol Ther 36, 467–476 (2012).

51. Chu, C. et al. Does pyogenic liver abscess increase the risk of delayed-onset primary liver cancer? Medicine (Baltimore). 96, (2017).

52. Lin, Y. et al. Seroepidemiology of *Klebsiella pneumoniae* colonizing the intestinal tract of healthy chinese and overseas chinese adults in Asian countries. BMC Microbiol 12, (2012).

53. Chung, D. R. et al. Fecal carriage of serotype K1 *Klebsiella pneumoniae* ST23 strains closely related to liver abscess isolates in Koreans living in Korea. Eur J Clin Microbiol Infect Dis 31, 481–486 (2012).

54. Qu, K. et al. Pyogenic liver abscesses associated with nonmetastatic colorectal cancers: An increasing problem in Eastern Asia. World J Gastroenterol 18, 2948–2955 (2012).

55. Jeong, S. W. et al. Cryptogenic pyogenic liver abscess as the herald of colon cancer. J Gastroenterol Hepatol 27, 248–255 (2012).

56. Garsin, D. A. Ethanolamine Utilization in Bacterial Pathogens: Roles and Regulation. Nat Rev Microbiol 8, 290–295 (2010).

57. Tsoy, O., Ravcheev, D. & Mushegian, A. Comparative Genomics of Ethanolamine Utilization. J Bac 191, 7157–7164 (2009).

58. Blin, C., Passet, V., Touchon, M., Rocha, E. P. C. & Brisse, S. Metabolic diversity of the emerging pathogenic lineages of *Klebsiella pneumoniae*. Environ. Microbiol. 19, 1881–1898 (2017).

59. Lin, Y. et al. Assessment of hypermucoviscosity as a virulence factor for experimental *Klebsiella pneumoniae* infections : comparative virulence analysis with hypermucoviscosity-negative strain. BMC Microbiol 11, (2011).

60. Su, S.-C., Siu, L. K., Yeh, K., Fung, C.-P. & Lin, J.-C. Community-Acquired Liver Abscess Caused by Serotype K1 *Klebsiella pneumoniae* with CTX-M-15-Type Extended-Spectrum β-Lactamase. Antimicrob Agents Chemother 52, 804–805 (2008).

61. Conlan, S. et al. Plasmid Dynamics in KPC-Positive *Klebsiella pneumoniae* during Long-Term Patient Colonization. MBio 7, e00742-16. (2016).

62. Stoesser, N. et al. Genome sequencing of an extended series of NDM-producing *Klebsiella pneumoniae* isolates from neonatal infections in a Nepali hospital characterizes the extent of community-Versus hospital-associated transmission in an endemic setting. Antimicrob. Agents Chemother. 58, 7347–7357 (2014).

63. Chung The, H. et al. A high-resolution genomic analysis of multidrug-resistant hospital outbreaks of *Klebsiella pneumoniae*. EMBO Mol. Med. 7, 227–39 (2015).

64. Wyres, K. L. & Holt, K. E. *Klebsiella pneumoniae* Population Genomics and Antimicrobial-Resistant Clones. Trend Microbiol 24, 944–956 (2016).

65. Ravin, A. W. Reciprocal Capsular Transformations of Pneumococci. JBac 77, 296–309 (1959).

66. Wyres, K. L. et al. Extensive Capsule Locus Variation and Large-Scale Genomic Recombination within the *Klebsiella pneumoniae* Clonal Group 258. Genome Biol. Evol. 7, 1267–1279 (2015).

67. Zhou, K. et al. Use of whole-genome sequencing to trace, control and characterize the regional expansion of producing ST15 *Klebsiella pneumoniae*. Sci Rep 6, 20840 (2016).

68. Lin, A. C. et al. Complete Genome Sequence of *Klebsiella pneumoniae* 1084, a Hypermucoviscosity-Negative K1 Clinical Strain. J Bac 194, 6316 (2012).

69. Gorrie, C. L. et al. Gastrointestinal carriage is a major reservoir of *K. pneumoniae* infection in intensive care patients. Clin Infect Dis cix270. do, (2017).

70. Davis, G. S. et al. Intermingled *Klebsiella pneumoniae* Populations between Retail Meats and Human Urinary Tract Infections. Clin Infect Dis 61, 892–899 (2015).

71. Follador, R. et al. The diversity of *Klebsiella pneumoniae* surface polysaccharides. Microb. Genomics 2, (2016).

72. Lin, H. et al. Two Genome Sequences of *Klebsiella pneumoniae* Strains with Sequence Type 23 and Capsular Serotype K1. Genome Announc. 4, e01097-16 (2016).

73. Wattam, A. R. et al. PATRIC, the bacterial bioinformatics database and analysis resource. Nucleic Acids Res 42, D581–D591 (2014).

74. Stoesser, N. et al. Predicting antimicrobial susceptibilities for *Escherichia coli* and *Klebsiella pneumoniae* isolates using whole genomic sequence data. J. Antimicrob. Chemother. 68, 2234–2244 (2013).

75. Wick, R. R., Judd, L. M., Gorrie, C. & Holt, K. E. Unicycler: Resolving bacterial genome assemblies from short and long sequencing reads. PLoS Comput Biol 13, e1005595 (2017).

76. Bankevich, A. et al. SPAdes: A New Genome Assembly Algorithm and Its Applications to Single-Cell Sequencing. J Comp Biol 19, 455–477 (2012).

77. Seemann, T. Prokka: Rapid prokaryotic genome annotation. Bioinformatics 30, 2068–2069 (2014).

78. Inouye, M. et al. SRST2: Rapid genomic surveillance for public health and hospital microbiology labs. Genome Med. 6, 90 (2014).

79. Gupta, S. K. et al. ARG-ANNOT, a New Bioinformatic Tool To Discover Antibiotic Resistance Genes in Bacterial Genomes. Antimicrob Agents Chemother 58, 212–220 (2014).

80. Carattoli, A. et al. In Silico Detection and Typing of Plasmids using PlasmidFinder and Plasmid Multilocus Sequence Typing. Antimicrob Agents Chemother 58, 3895–3903 (2014).

81. Rutherford, K. et al. Artemis : sequence visualization and annotation. Bioinformatics 16, 944–945 (2000).

82. Wick, R. R., Schultz, M. B., Zobel, J. & Holt, K. E. Bandage: interactive visualization of de novo genome assemblies. 31, 3350–3352 (2015).

83. Croucher, N. J. et al. Rapid phylogenetic analysis of large samples of recombinant bacterial whole genome sequences using Gubbins. Nucleic Acids Res. 43, e15 (2015).

84. Stamatakis, A. RAxML-VI-HPC: Maximum likelihood-based phylogenetic analyses with thousands of taxa and mixed models. Bioinformatics 22, 2688–2690 (2006).

85. Ashkenazy, H. et al. FastML: a web server for probabilistic reconstruction of ancestral sequences. Nucleic Acids Res 40, W580–W584 (2012).

86. Rambaut, A., Lam, T. T., Carvalho, L. M. & Pybus, O. G. Exploring the temporal structure of heterochronous sequences using TempEst (formerly Path-O-Gen). Virus Evol. 2, vew007 (2016).

87. To, T.-H., Jung, M., Lycett, S. & Gascuel, O. Fast Dating Using Least-Squares Criteria and Algorithms. Syst Biol 65, 82–97 (2016).

88. Drummond, A. J., Suchard, M. A., Xie, D. & Rambaut, A. Bayesian Phylogenetics with BEAUti and the BEAST 1.7. Mol Biol Evol 29, 1969–1973 (2012).

89. Darling, A. C. E., Mau, B., Blattner, F. R. & Perna, N. T. Mauve : Multiple Alignment of Conserved Genomic Sequence With Rearrangements. Methods 14, 1394–1403 (2004).

90. Zhou, Y., Liang, Y., Lynch, K. H., Dennis, J. J. & Wishart, D. S. PHAST : A Fast Phage Search Tool. Nucleic Acids Res 39, W347–W352 (2011).

91. Page, A. J. et al. Roary: rapid large-scale prokaryote pan genome analysis. Bioinformatics 31, 3691–3693 (2015).

92. Bland, C. et al. CRISPR Recognition Tool (CRT): a tool for automatic detection of clustered regularly interspaced palindromic repeats. BMC Bioinformatics 8, 1–8 (2007).

93. Li, W. & Godzik, A. Cd-hit: a fast program for clustering and comparing large sets of protein or nucleotide sequences. Bioinformatics 22, 1658–1659 (2006).

